# A multiplexed plant-animal SNP array for selective breeding and species conservation applications

**DOI:** 10.1101/2022.09.07.507051

**Authors:** Sara Montanari, Cecilia Deng, Emily Koot, Nahla V. Bassil, Jason D. Zurn, Peter Morrison-Whittle, Margaret L. Worthington, Rishi Aryal, Hamid Ashrafi, Julien Pradelles, Maren Wellenreuther, David Chagné

## Abstract

Reliable and high-throughput genotyping platforms are of immense importance for identifying and dissecting genomic regions controlling important phenotypes, supporting selection processes in breeding programmes, and managing wild populations and germplasm collections. Amongst available genotyping tools, SNP arrays have been shown to be comparatively easy to use and generate highly accurate genotypic data. Single species arrays are the most commonly used type so far; however, some multi-species arrays have been developed for closely related species that share SNP markers, exploiting inter-species cross-amplification. In this study, the suitability of a multiplexed plant-animal SNP array, including both closely and distantly related species, was explored. The performance of the SNP array across species for diverse applications, ranging from intra-species diversity assessments to parentage analysis, was assessed. Moreover, the value of genotyping pooled DNA of distantly related species on the SNP array as a technique to further reduce costs was evaluated. SNP performance was generally high, and species-specific SNPs proved suitable for diverse applications. The multi-species SNP-array approach reported here could be transferred to other species to achieve cost savings resulting from the increased throughput when several projects use the same array, and the pooling technique adds another highly promising advancement to additionally decrease genotyping costs by half.

## Introduction

The development of medium-density genotyping tools for the inexpensive, rapid and reliable screening of hundreds of samples is of utmost importance in molecular breeding programmes and for the management of wild populations and germplasm collections. Nowadays, there are a number of high-throughput genotyping technologies and platforms, each with advantages and disadvantages. Whole genome sequencing (WGS) is increasingly being used; however, it can be a costly approach for the routine screening of large numbers of samples, particularly in species with large genome sizes. While reduced-representation genotyping-by-sequencing (GBS [1]; restriction-site associated DNA sequencing (RADseq) [2]) is more scalable, such methods generate high error rates and missing data [3]. Additionally, extensive bioinformatics resources are required for the curation of GBS datasets [4]. Single nucleotide polymorphism (SNP) arrays generate genotype data that are more reliable than GBS [5,6], and their costs can be relatively low, particularly in comparison with WGS approaches. One limitation of SNP arrays is that the identification of variants and the design of efficient SNP probes depend on the availability of a well-assembled low-error rate reference genome. However, this requirement has become less of a problem recently, as high-quality genomes have been developed for many commercially and ecologically relevant species [7,8]. The main disadvantage of SNP arrays is the ascertainment bias caused by over-representing diversity in the re-sequencing panels during the polymorphism detection step. However, that is not a major concern if SNP arrays are applied to screen populations that are genetically related to the re-sequencing panels. SNP arrays have proven particularly useful for genome-informed breeding applications, such as checking sample identity, pedigree and relationship assignment, trait mapping and genomic predictions [9–14]. Nonetheless, when applying SNP arrays to screen wild/natural populations, increased attention needs to be given to the possibility that genetic variation can be missed. This is particularly likely in cases where the re-sequencing panels used for polymorphism detection are small and/or are not collected across the same spatial scale as subsequently tested samples. Nevertheless, SNP arrays can be powerful tools for wild population analyses, provided the polymorphism detection panels are large and diverse, and they can address questions related to population structure, provenance, and kinship.

A large number of SNP arrays have been designed for plant and animal species important for primary production (e.g. agriculture, horticulture, aquaculture), with densities ranging from few hundred to several thousand markers ([5,6,15–21] to cite some of the most recent). In some cases, researchers have exploited the known synteny among genomes of sister or closely related species and developed SNP arrays that combine markers from different genera. Key examples are the apple (*Malus domestica*) and pear (*Pyrus communis*) Illumina Infinium II 9K SNP array [22], the Pacific (*Crassostrea gigas*) and European oysters (*Ostrea edulis*) Axiom™ 60K SNP array [23], the “MedFish” Axiom™ 60K SNP array for European seabass (*Dicentrarchus labrax*) and gilthead seabream (*Sparus aurata*) [17], and the Axiom™ SerraSNP array with ∼30K SNPs each for the South American fresh water fishes pacu (*Piaractus mesopotamicus*) and tambaqui (*Colossoma macropomum*) [16]. However, multi-species SNP arrays designed for different and completely unrelated taxa with the purpose of pooling DNA from two (or more) samples have not been explored thus far. The development and validation of such an array was the objective of this work, in an effort to create a resource for routine, medium-density genotyping of hundreds of samples from breeding programmes as well as wild populations for four genera with rapidly growing primary production industries. These included two plant and two fish genera, specifically *Rubus, Leptospermum, Chrysophrys* and *Pseudocaranx*.

The *Rubus* genus of the Rosaceae family comprises red (*R. idaeus*) and black (*R. occidentalis*) raspberries, as well as blackberries (*Rubus* subgenus *Rubus*). Simple sequence repeat (SSR) markers were used to confirm identity and assess diversity at the Agricultural Research Service of United States Department of Agriculture (USDA-ARS) - National Clonal Germplasm Repository (NCGR) [24,25], and target capture sequencing was used for phylogenetic analyses [26]. However, while genetic maps and reference genomes are now available for *Rubus* [27–31], the application of genomics resources in these crops is still limited [32]. Similarly, the shrub *L. scoparium*, which includes New Zealand mānuka, is an undomesticated species supporting the production of honeys with unique high antimicrobial properties which attract high premium prices. The recent publication of a reference genome for *L. scoparium* [33] and the re-sequencing of specimens from across its natural range [34] have opened new avenues for the development of genomic resources for this species to support its management and selective breeding. Emerging animal species of economic or ecological importance would also benefit from an increased application of genomic tools. Australasian snapper (*C. auratus*, tāmure) and silver trevally (*P. georgianus*, araara) are two candidate species for aquaculture in New Zealand. Selective breeding programmes for both these finfish species have recently been initiated, to diversify the aquaculture sector [35–37]. Mānuka, tāmure and araara are native to New Zealand (Aotearoa) and considered treasures (taonga) by Māori [38], who have traditional uses for them as sources of food and medicine.

This study describes the design of a multi-species plant-animal 60K SNP array that combines 13K SNP markers for *Rubus*, 9K for mānuka, 18K for snapper and 20K for trevally, and its validation by the screening of more than 6,000 pooled plant/fish DNA samples. The potential of DNA pooling as a strategy to reduce genotyping costs was also evaluated. Pedigree reconstruction and genetic diversity analyses were performed to demonstrate the use of the SNP array. Finally, cross-amplification in species closely related to snapper (Japanese red seabream, *Chrysophrys/Pagrus major*) and silver trevally (yellowtail kingfish, *Seriola lalandi*) was examined.

## Materials and Methods

### Sequencing, variant calling and SNP filtering

Different sequencing datasets were available for each of the four organisms targeted in this study and the variant calling and filtering analyses performed are described in S1 Appendix (*Rubus* spp.), S2 Appendix (mānuka), S3 Appendix (snapper) and S4 Appendix (trevally).

### SNP final selection and array design

For the selected SNPs for each of the species, 60 bp up- and down-stream flanking sequences were extracted from the respective reference genomes and submitted to Thermo Fisher Scientific for quality scoring. Only SNPs for which at least one probe was recommended (pconvert > 0.6, no wobbles, and poly count = 0) were kept. BLASTn analysis was then performed to remove plant SNPs with flanking sequences that showed high similarity to fish DNA, and *vice versa*, using the reference genomes for all four species, to avoid cross-hybridization between fish and plant DNA. Finally, SNPs that were too close to the start of the chromosome/scaffold (< 60 bp), and for which flanking sequences could not be extracted, were eliminated.

For the raspberry SNPs, alignments to a *R. idaeus* ‘Heritage’ draft genome (Driscoll’s, Watsonville, CA, USA; unpublished) were checked, and those that had either no hits or more than one mismatch were discarded. This quality check was necessary because the SNPs were designed on the *R. occidentalis* reference genome, the only raspberry genome publicly available at this time, while target populations for application of this SNP array are mainly derived from *R. idaeus*.

Finally, probes for all selected SNPs were tiled on an Applied Biosystems® Axiom™ myDesign™ array (Thermo Fisher Scientific).

### DNA extractions

SNP array genotyping reactions were performed by multiplexing one fish and one plant DNA sample in equal quantities. *Rubus* and mānuka leaf samples were collected by a consortium of different institutes and DNA extracted in the respective laboratories. *Rubus* DNA extractions were performed in 96-well plates from freeze-dried tissue using commercial kits (e.g. Macherey-Nagel Nucleospin kit was used at PFR). Mānuka samples were collected in natural stands in remote locations using silica beads in 2mL screw cap tubes as described in Koot *et al*. [34] and DNA was extracted using a modified CTAB protocol [39,40]. Fish DNA extractions were performed using fin clip samples collected in 96-well plates and using a proprietary automated protocol by Slipstream Automation Ltd (Palmerston North, New Zealand). All fish and plant DNA samples were then dried for shipping to Labogena (Jouy-en-Josas, France), where they were quantified using fluorometry, normalised to 2*µ*g and mixed.

### SNP array genotyping

Two separate batches of genotyping were performed, amounting to 2,686 and 3,455 samples, respectively, and including both pooled plant/fish DNA samples and single-species ones. Specifically, these were: 49 *Rubus* only samples, 511 mānuka only, 214 Australasian snapper only, 322 silver trevally only, 2,908 *Rubus* + snapper pooled samples, 39 *Rubus* + Japanese red seabream, 225 *Rubus* + trevally, 23 *Rubus* + kingfish, 1,122 mānuka + snapper, and 656 mānuka + trevally (S3 Table). In total, 3,244 *Rubus*, 2,289 mānuka, 4,244 snapper, 1,203 trevally, 39 red seabream, and 23 kingfish samples were genotyped.

The SNP data were automatically split for the four sets of SNPs (*Rubus*, mānuka, snapper and trevally) and analysed separately in the Axiom Analysis Suite v5.1.1 software (https://www.thermofisher.com/nz/en/home/life-science/microarray-analysis/microarray-analysis-instruments-software-services/microarray-analysis-software/axiom-analysis-suite.html). The data were quality-filtered using a Dish QC (DQC) threshold of 0.82 (default) and a QC call rate threshold of 95, and genotypes were called with the default parameters. The OTV caller was run on the *OffTargetVariants* (OTV), i.e. SNPs that might contain null alleles.

### SNP validation

The effectiveness of the SNP array was evaluated by verifying if the expected population structure could be depicted in subsets of samples for all four organisms. Only the higher quality SNPs (*PolyHighResolution* SNPs (PHR) and *NoMinorHom* (NMH)) were used for these analyses. The methods applied in each species are described in S1 Appendix (*Rubus* spp.), S2 Appendix (mānuka), S3 Appendix (snapper and seabream) and S4 Appendix (trevally).

### Comparison of genotyping quality among species

The quality parameters DQC and QC call rate were compared among species using boxplots. The effect of DNA pooling on the quality of genotyping was also assessed for each of the four main species by generating boxplots, calculating descriptive statistics such as mean, standard deviation and median, and running an Unequal Variance (Independent) T-test (or Welch T-test). Correlations between quality values of plant and fish samples from the same reaction were depicted in scatter plots. Finally, correlations between DQC and QC call rate values for the four main species were observed using scatter plots. All samples submitted for genotyping were used for these comparisons.

### Ethics statement

Informed consent was granted verbally by Māori landowners to re-use the reference genome of *L. scoparium* ‘Crimson Glory’ [33], as well as the pool-sequencing data and DNA samples of Koot *et al*. [34], for the purpose of developing and evaluating the SNP array in mānuka. The consent was given during a project meeting in June 2020 and minutes were documented. All trevally research carried out in this study was reviewed and approved by the animal ethics committee of Victoria University of Wellington in New Zealand (Application number 25976). For snapper, ethics approval was granted through Victoria University of Wellington in New Zealand (Application number 2014R19) and the University of Auckland (Ref. 002169).

## Results

### Multi-species SNP array design

#### *Rubus* spp

##### Raspberry

Variants were called from GBS datasets of seven F_1_ *Rubus* subgenus Idaeobatus populations (S1 Table). The total numbers of variants called from the populations X14.102, X16.015 and NC493×CW datasets were, respectively, 1,335,709, 406,253, and 649,597, which were then reduced in turn to 432,433, 239,830, and 403,211 SNPs after initial basic filtering; 26, one, and 10 samples respectively with high rate of missing data were removed from each population. For the dataset including families X16.093, X16.095, X16.109, and X16.111, a total number of 53,745 variants were called across 358 samples, and filtered down to 52,694 SNPs. Merging of VCF files from all seven populations and thinning resulted in 398,596 unique SNPs with no other neighbouring polymorphism. Finally, after extraction of the “validated SNPs” (i.e. SNPS that successfully grouped into Linkage Groups (LGs) in JoinMap v5.0 [41]) and removal of the A/T and C/G SNPs, this number reduced to 25,910, including the 859 SNPs associated with sugar content [42]. Of these, 23,898 SNPs had at least one probe recommended, and the 9,376 of them that aligned well to the *R. idaeus* ‘Heritage’ genome were included in the array.

##### Blackberry

A total of 4,163 SNPs were selected from a WGS dataset of 27 blackberry accessions, including 1,864 SNPs evenly distributed throughout the genome and 2,299 SNPs within genes of interest potentially associated with sweetness [42], thornlessness [43], and flowering [31]. Of these selected SNPs, 3,719 aligned to a unique position in the *R. occidentalis* reference genome and were submitted to Thermo Fisher Scientific for scoring. The 3,347 loci with recommended probes were included in the final array.

### Mānuka

Quality filtering of the mānuka pooled sequencing datasets from Koot *et al*. [34] resulted in 2,006,036 SNPs. Eight classes of SNPs were identified based on their MAF values in each or a combination of gene pools (Northern North Island (NNI), the Central and Southern North Island (CNI), the East Coast North Island (ECNI), and two gene pools in the South Island representing the North-East (NESI) and South-West of the South Island (SWSI); S6 Table), and a total of 42,122 SNPs were identified that fitted into these classes. Thermo Fisher Scientific scoring classified 33,484 of these SNPs as recommended, and a random set of 9,002 SNPs evenly distributed in the genome was selected for inclusion in the array.

### Snapper

A total of 6,255,825 SNPs resulted from variant calling on 80 re-sequenced samples from the PFR snapper breeding programme, reduced to 4,151,564 after thinning with a 30bp window. No SNP site exhibited > 20% missing data, and filtering for maximum DP, MAF, multi-allelic and A/T and C/G SNPs resulted in a total of 2,419,846 SNPs. After LD pruning, 26,719 SNPs were left, of which 13 were removed because they either aligned to the raspberry or mānuka genomes, or because they were too close to the terminal of the scaffold. Finally, 22,238 SNPs were recommended by Thermo Fisher Scientific scoring, and 18,489 were included in the array. It was noted that 13,204 of these SNPs were in coding regions, according to the male and female gene annotation for the *C. auratus* v 1.0 reference genome [44].

### Trevally

The initial SNP calling performed by Valenza-Troubat *et al*. from WGS reads of 13 trevally samples [37] resulted in a dataset of 17,795,808 SNPs, which were then thinned to 7,969,343 and subsequently reduced to 3,087,247 after filtering for missing data, maximum DP, MAF, multi-allelic and A/T ad C/G SNPs. Afterwards, LD pruning left 26,666 SNPs and 264 had to be removed because of risk of cross-hybridization with plant genomes or because they were too close to a scaffold terminal. Thermo Fisher Scientific scoring recommended 22,076 SNPs, and 20,234 were successfully included in the array.

In summary, the Axiom multi-species plant-animal 60K SNP array includes 60,448 SNPs from five species and four genera: *R. idaeus, R*. subgenus *Rubus, L. scoparium, C. auratus* and *P. georgianus* (Table 1, S7 Table).

**Table 1.** Number of single nucleotide polymorphism (SNP) markers for each species in the Axiom multi-species 60K SNP array.

| Organism | Species | # SNP markers |
| --- | --- | --- |
| Raspberry | *Rubus idaeus/occidentalis* | 9,376 |
| Blackberry | *Rubus* subgenus *Rubus* | 3,347 |
| Total *Rubus* spp. |  | 12,723 |
| Mānuka | *Leptospermum scoparium* | 9,002 |
| Snapper | *Chrysophrys auratus* | 18,489 |
| Trevally | *Pseudocaranx georgianus* | 20,234 |
| Total |  | 60,448 |

### Genotyping of test sample sets for each species and validation of the SNP array

#### *Rubus* spp

##### Diploids

Of the 477 diploid samples analysed in this study, 57 and 82 failed to pass the DQC and the QC call rate thresholds, respectively, leaving a set of 338 samples for further statistical analyses. Cluster plot assessment highlighted a subset of samples that frequently fell out of the posterior cluster margins (S2 Figure), causing errors in the automatic quality evaluation and classification of the SNP markers. A threshold of 0.85 for “allele_deviation_mean” was then established to remove the low-quality samples, leaving 305 samples to be re-analysed. Finally, 276 samples from PFR germplasm and 29 from the NCGR passed all quality thresholds and exhibited an average call rate of 99.0%. Genotyping resulted in 6,141 SNPs classified as PHR, 1,211 as NMH, 612 as OTV, 1,669 as *MonoHighResolution*, 2,622 as Other and 468 as *CallRateBelowThreshold* (S8 Table). Two or more replicates were successfully genotyped for four different accessions, including BC 64-9-81 (two replicates), ‘Glen Ample’ (two replicates), ‘Wakefield’ (five replicates), and *R. spectabilis* ‘Gibbs Lake’ (three replicates). All replicates had identity-by-state (IBS) > 0.97 except for ‘Wakefield’, which grouped into two sets of duplicated samples each with IBS > 0.97. Two of the ‘Wakefield’ samples that were identical to each other but different from the other three were later ascertained to be sampling errors. However, since they were technical replicates from the same sample, they were included in the analysis. Over all the PHR and NMH SNPs verified, 6,885 (93.7%) had no genotyping inconsistencies and were considered robust. Of these, 6,235 were raspberry SNPs and 650 blackberry SNPs. Principal Component Analysis (PCA) showed three main clusters along the PC1 (25.4% of variation explained), with almost all samples from the NCGR grouping together into one cluster (Fig 1A). However, there was no correspondence between the PCA clustering and the species assignment (Fig 1B). A Discriminant Analysis of Principal Components (DAPC) was run using the optimal number of 11 clusters, 100 PCs and 4 DAs. A three-dimensional scatter plot showed a large group that included seven of the 11 clusters, and four small well-separated groups (Figs 1C and D). There was good correspondence between the three main clusters identified with the PCA and the three-dimensional separation observed with the DAPC (Fig 1C). However, the DAPC clusters still could not be explained by the taxonomy of the samples (Fig 1D).

**Fig 1.**
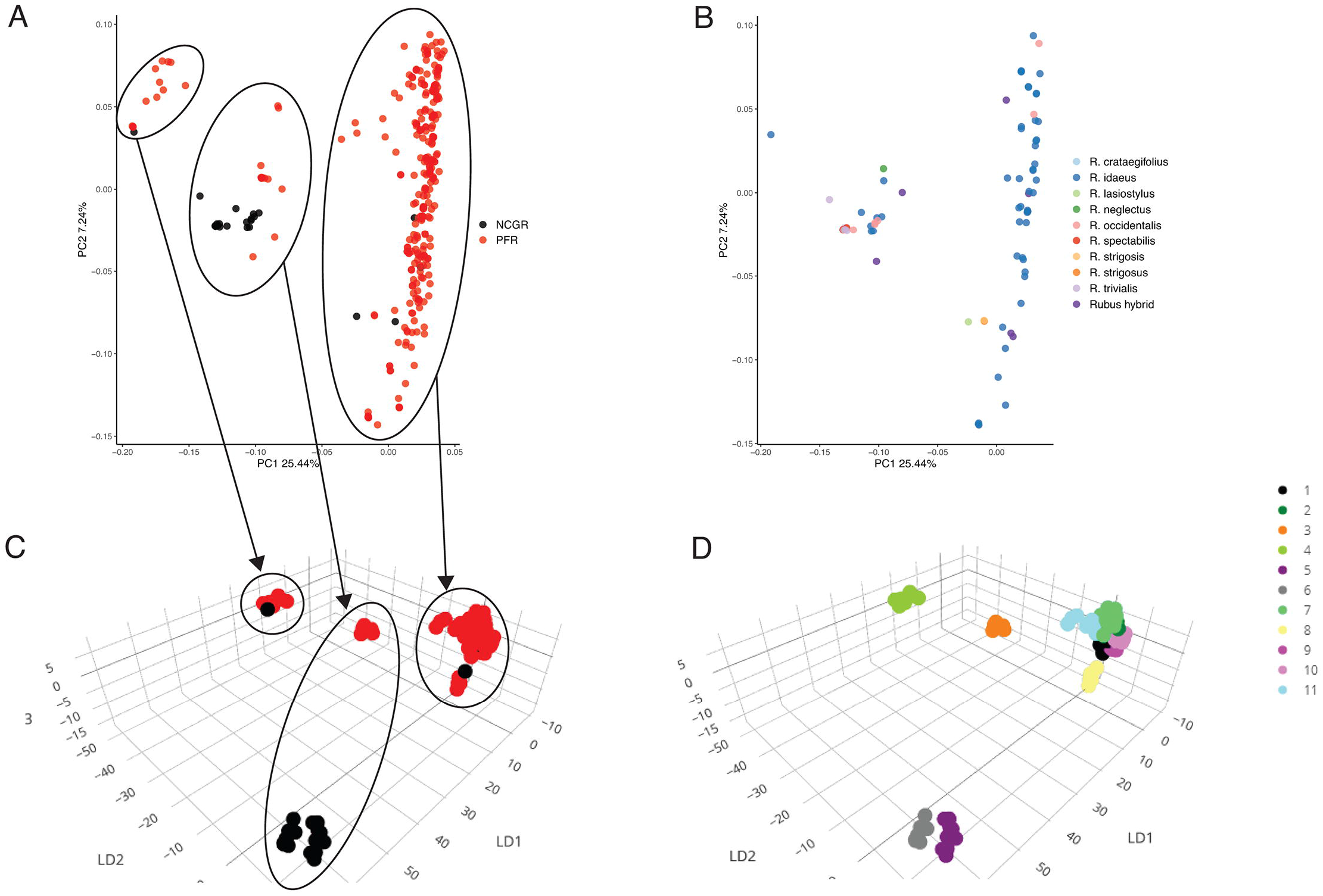
Genetic diversity of diploid *Rubus* samples. Principal Component Analysis with samples coloured by repository (A) and assigned species (B). Discriminant Analysis of Principal Components plot with samples coloured according to repository (C) and DAPC clustering (D). Circles and arrows in plots A and C show correspondence between PCA and DAPC clustering. NCGR = USDA-ARS National Clonal Germplasm Repository; PFR = The New Zealand Institute for Plant and Food Research Ltd.

##### Tetraploids

Tetraploid models were successfully fitted in fitPoly v3.0.0 [45,46] for 4,872 SNP markers, out of the total 12,723, in 739 samples. This dataset was further reduced to 666 samples and 4,388 SNPs after filtering for missing rate. These included 2,899 raspberry and 1,489 blackberry SNPs. Three main clusters could be observed on the PC1 (27.1% of variation explained) versus PC2 (7%) plot, with two clusters very close together, and with several samples remaining ungrouped (Fig 2). As for the diploid samples, the results of the PCA could not be explained by the reported taxonomy; however, there was good correspondence with the collection of origin. Most of the PFR samples formed two of the main clusters, while some overlapped with the third cluster that mainly contained UArk samples; ungrouped samples were mostly from NCGR. Samples from PFR and UArk included parental and seedling selections from their breeding programmes, while accessions from NCGR represented a diverse range of species and genotypes maintained at their repository for conservation purposes.

**Fig 2.**
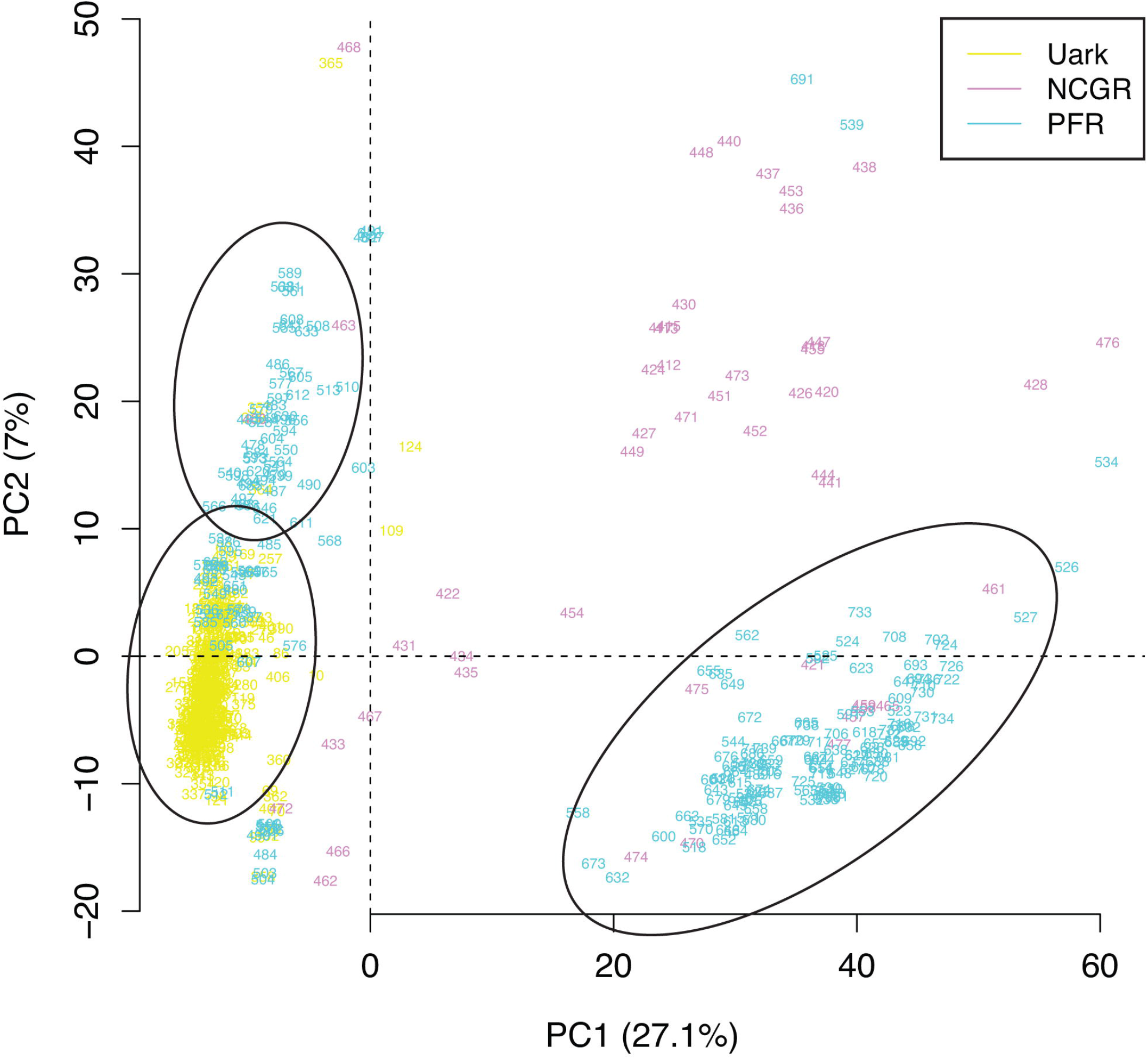
Genetic diversity of tetraploid *Rubus* samples. Principal Component Analysis with samples coloured by repository. Uark = University of Arkansas System Division of Agriculture; NCGR = USDA-ARS National Clonal Germplasm Repository; PFR = The New Zealand Institute for Plant and Food Research Ltd.

### Mānuka

Of the 264 mānuka samples screened using the array, 233 passed the QC filters. Of the 31 failed samples, seven were because of a low DQC score and 24 were because of a QC call rate < 95. Of the 9,002 SNPs included in the array, 4,969 were classified as PHR, 1,026 as OTV and 124 as NMH. These 6,119 polymorphic good quality markers represented 68% of the mānuka SNPs included in the array. Of the 2,883 unsuccessful SNPs, 19 were monomorphic (*MonoHighResolution*), 761 had call rate below the threshold and the majority (2,103) had poor clustering (S8 Table). When K-means clustering and DAPC analyses were performed using only the 4,969 PHR SNPs, the 233 samples separated into four clusters matching four geographical regions: NNI, CSNI, ECNI and South Island (SI) (Fig 3A; S4 Table). *F*_*ST*_ calculated between each of the four regions ranged from 0.08 between ECNI and CSNI, to 0.20 between ECNI and NNI (Fig 3B).

**Fig 3.**
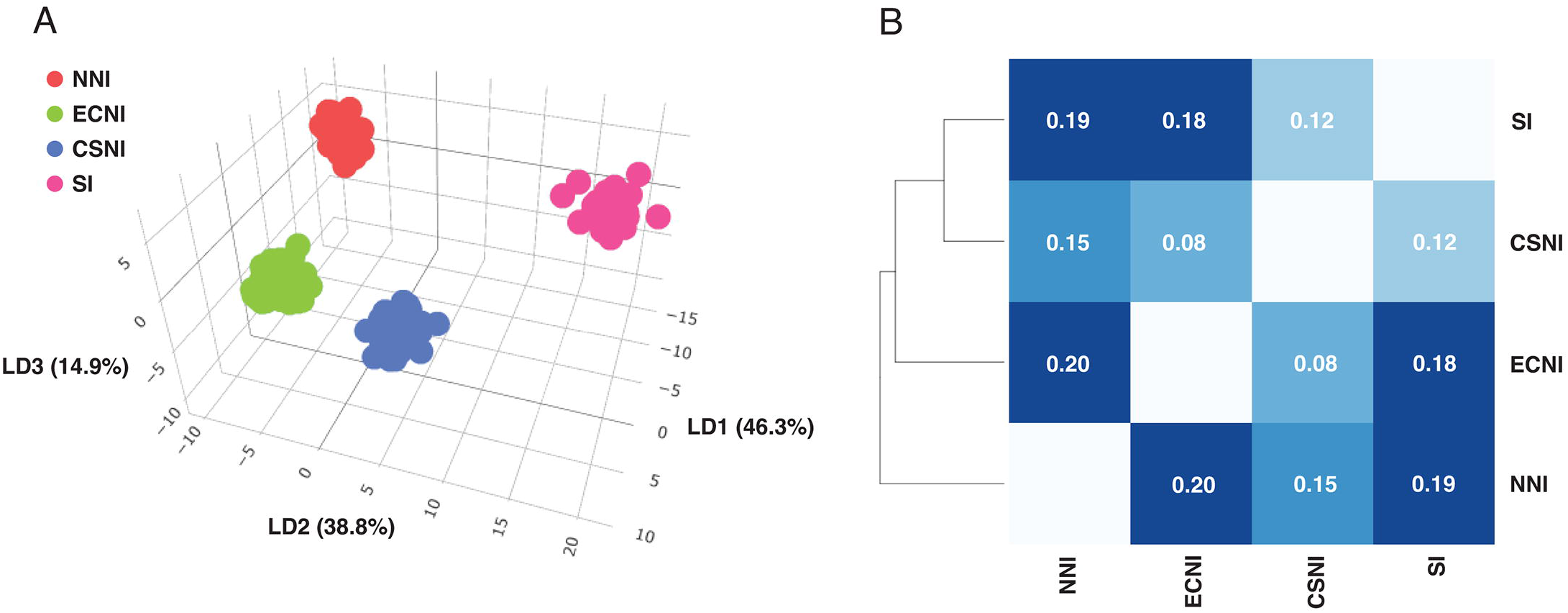
Population genetics analysis of mānuka samples from New Zealand. (A) Discriminant Analysis of Principal Components plot with samples coloured according to provenance. (B) *F*_*ST*_ analysis. NNI = Northern North Island; ECNI = East Cape North Island; CSNI = Central & Southern North Island; SI = South Island.

### Snapper and seabream

Of the 4,244 snapper samples screened using the array, 3,915 passed the QC filters. In the first batch, 48 samples failed because of a low DQC score and 121 because of a call rate < 95. In the second batch, 23 and 137 samples failed at DQC and QC call rate, respectively. The seabream samples were analysed together with the snapper samples from the first batch and only four out of 39 failed because of low QC call rate. Of the 18,489 SNPs included in the array, in the first batch 11,921 (64.5%) were classified as PHR and 941 (5.1%) as NMH, while the other were either monomorphic, OTV or were of poor quality. In the second batch numbers were similar, with 11,601 (62.8%) PHR and 1,000 (5.4%) NMH. 10,692 and 472 SNPs were classified as PHR and NMH in both batches, respectively (S8 Table), and were used for subsequent analysis. The variation explained by the PC1 and PC2 was 7.87% and 6.56%, respectively. Three major clusters could be observed, corresponding to the snapper broodstock of origin (Fig 4A). Examination of the PC3 vs PC4 plot (explaining variation of 3.81% and 2.73%, respectively) revealed a major central cluster including Broodstocks 2 and 3, while snapper samples from Broodstock 1 formed several separate clusters around it (Fig 4B). The seabream samples overlapped with the snapper Broodstock 1 in the PC1 vs PC2 plot, while they formed a distinct cluster in the PC3 vs PC4 plot (Fig 4B). Broodstock 1 included adult snapper individuals harvested from the wild and their direct offspring spawned in captivity, while Broodstocks 2 and 3 included adult individuals generated through PFR’s selective breeding programme as well as their direct offspring, which were also spawned in captivity. In total, 2,937 trios were detected and confirmed by Mendel test, which resulted in 90.9% of all offspring having assigned parentage (Fig 4C). Differences were observed among broodstock lines, with 98.3%, 95.5% and 79.8% of offspring assigned in Broodstocks 1, 2 and 3, respectively. In addition, the proportion of adults that contributed to the next generation differed substantially among broodstocks – with 51.7%, 14.3%, and 70.4% of adult fish contributing to offspring generation in Broodstocks 1, 2, and 3 respectively (Fig 4C).

**Fig 4.**
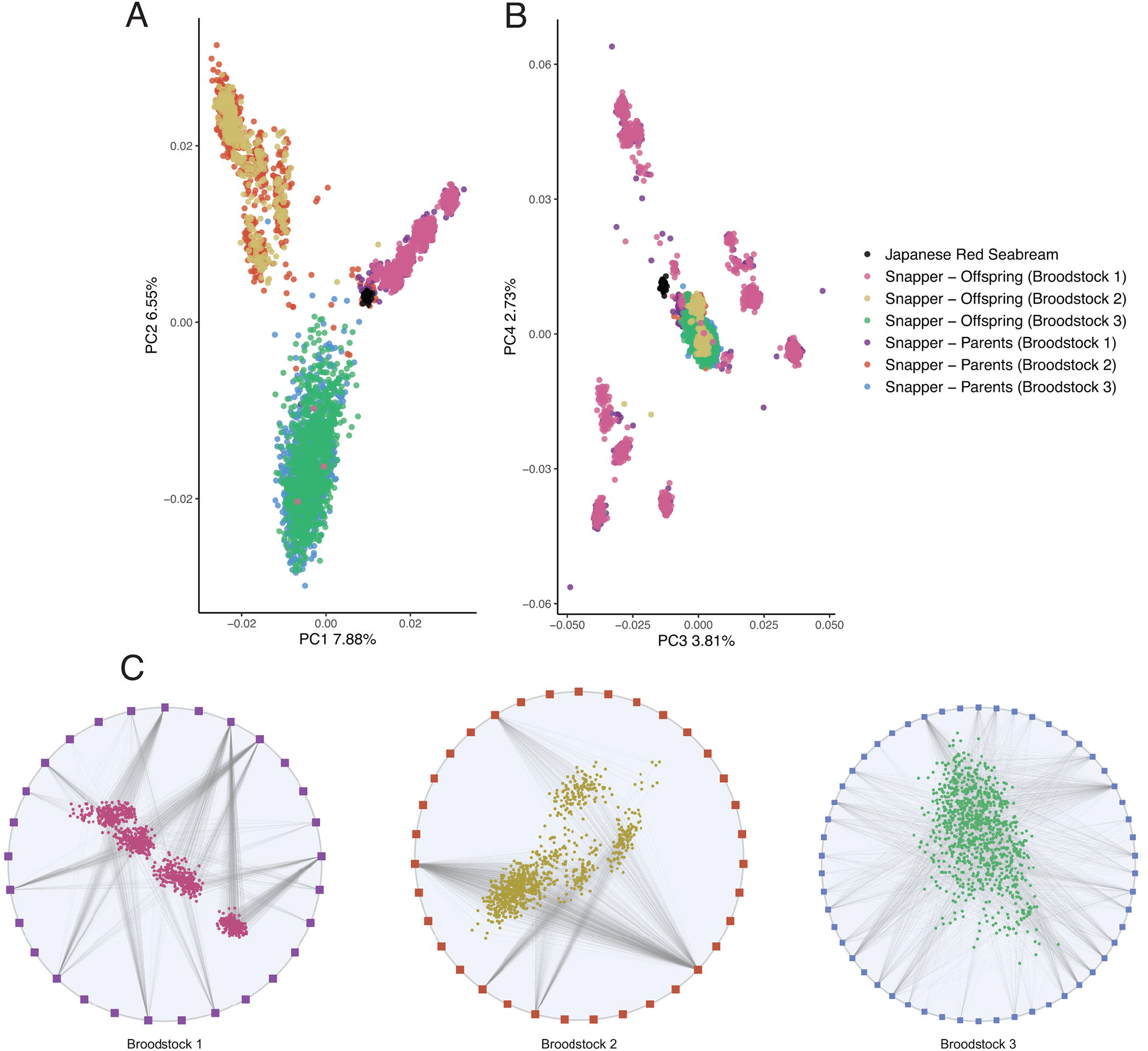
Population genetics analysis and pedigree reconstruction of Australasian snapper and Japanese red seabream. Principal Component Analysis coloured by species and population: PC1 vs PC2 (A) and PC3 vs PC4 (B). (C) Pedigree networks: parent-offspring relationships are indicated by a line connecting adult snapper (square points) and juvenile snapper (circular points) of the three broodstock lines.

### Trevally and kingfish

Since only 48 samples were submitted in the first batch, genotypes for these were called together with all trevally samples from the second batch, and 1,072 out of 1,203 passed all QC filters. Only 10 samples failed the DQC filter, and 121 did not pass the QC call rate threshold of 95. All kingfish samples, which were analysed together with the trevally, failed at DQC. Of the 20,234 SNPs included in the array, 10,157 were classified as PHR, 1,556 as NMH, 2,521 as OTV, 146 were monomorphic, 1,100 had call rate below the threshold, and 4,754 exhibited poor clustering (S8 Table). PHR and NMH SNPs were used for subsequent analysis. Of the 978 samples examined for SNP validation, 938 passed the Axiom QC genotyping filters; these included 54 individuals from Australia and 884 from New Zealand (S5 Table). The PCA showed a clear separation between Australian and New Zealand samples along the PC1, which accounted for 3.30% of the variation (Fig 5). *F*_*ST*_ between the two populations was estimated to be 0.19.

**Fig 5.**
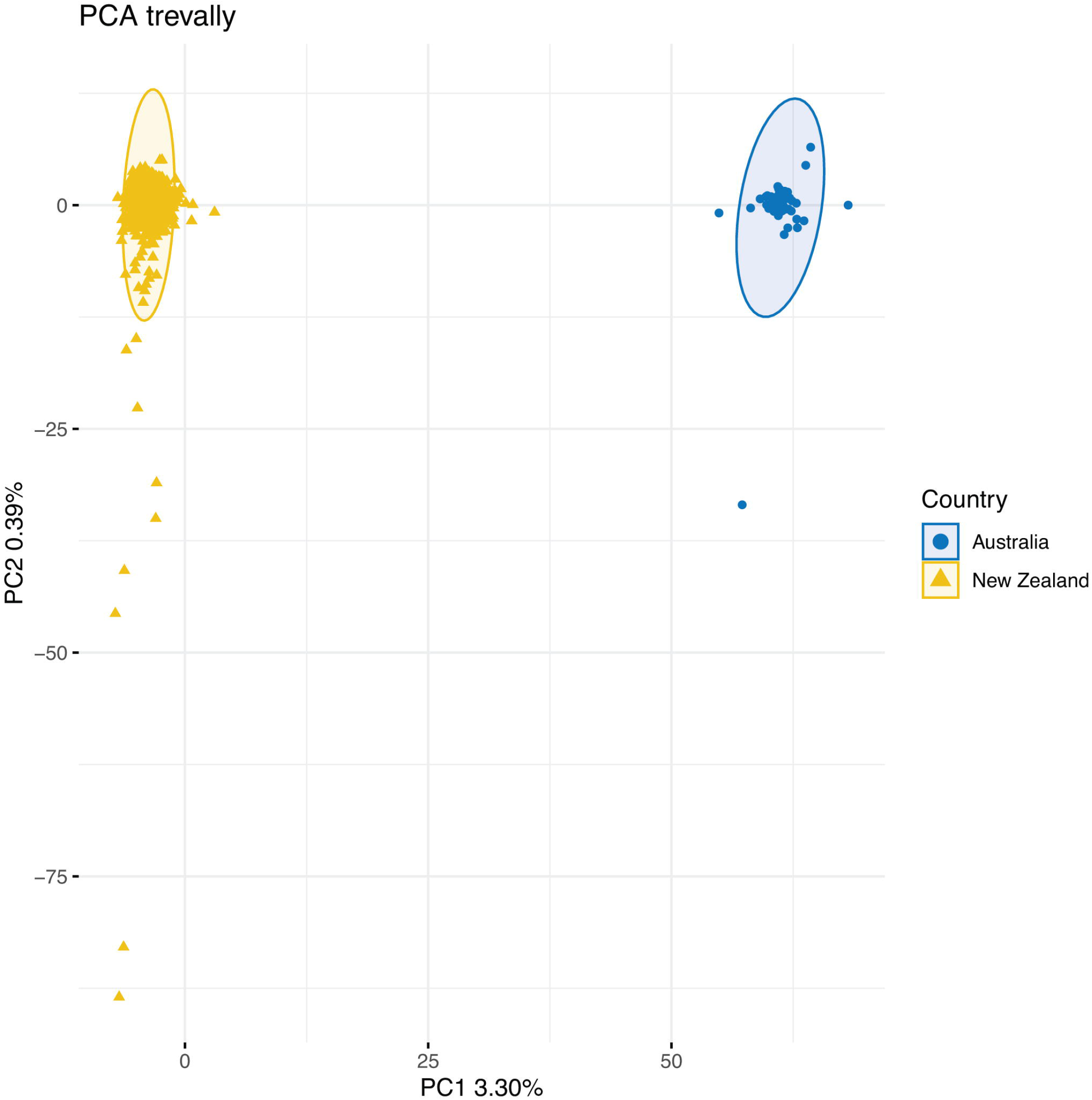
Principal Component Analysis of trevally samples caught in Australia and New Zealand.

### Evaluation of quality of genotyping

DQC values were always higher in fish samples than in plants, with the exception of kingfish; however, a high variability was observed for all species (Fig 6A). QC call rate was fairly uniform across all species (Fig 6B). In both plants and fishes, pooling appeared to have significant negative effect on the DQC (T-test, p-value < 0.05); however, differences were more marked in the plant than in the fish samples (Fig 7A). QC call rate appeared higher in the pooled plant samples than in the non-pooled ones, but SD was also larger, while the difference was not significant in fish (Fig 7B). Finally, a large number of multiplexed reactions that returned high DQC values for the fish samples exhibited low DQC for the corresponding plant ones (Fig 7C), while QC call rate values were overall in greater agreement between the two (Fig 7D).

**Fig 6.**
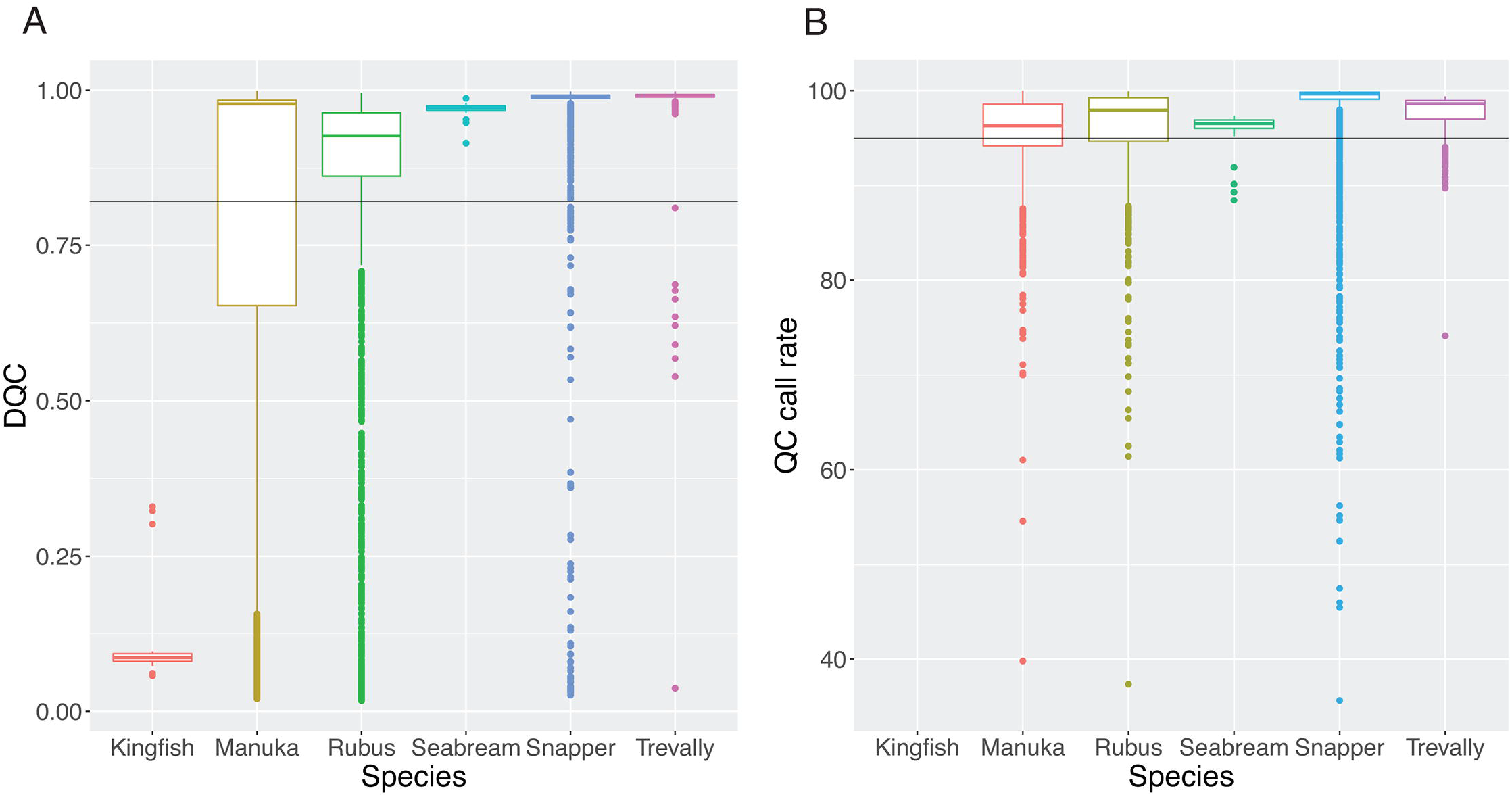
Comparison of quality parameters among species. Boxplots for Dish QC (DQC) (A) and QC call rate values (B) grouped by species.

**Fig 7.**
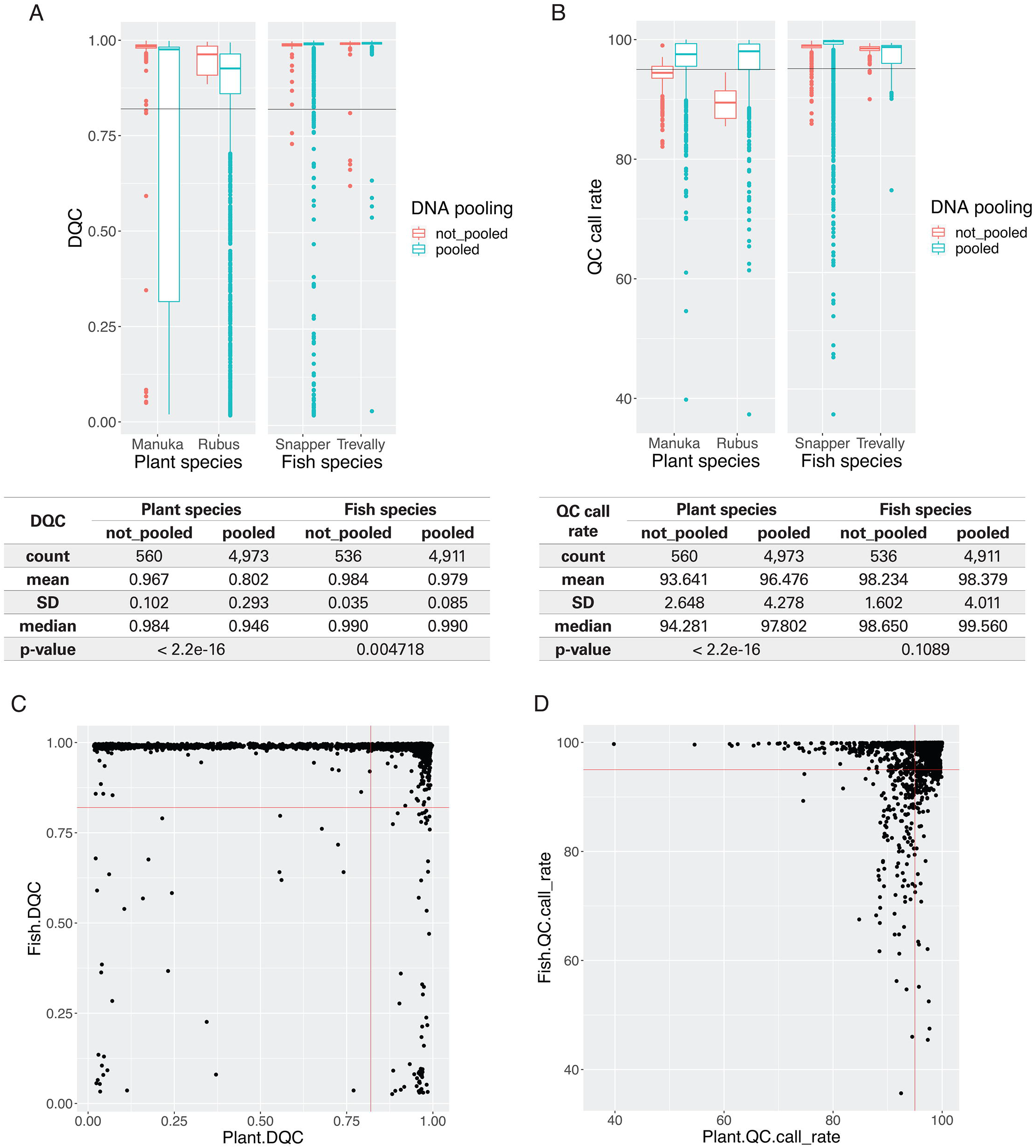
Comparison of quality parameters between pooled ad non-pooled DNA samples. Boxplots for Dish QC (DQC) (A) and QC call rate values (B) grouped by species and pooling, with corresponding values for sample count, mean, standard deviations (SD) and median. Scatter plot for DQC and QC call rate values of pooled plant versus fish samples (C and D).

## Discussion

Our work demonstrates the successful development of a multi-species plant-animal SNP array and the application of this array in selective breeding programmes and the management of natural populations and plant germplasm.

### Novel SNP array developed for two plant and two fish species

This is not only the first multi-species plant-animal SNP array, but also the first array for each of the included organisms *Rubus* spp., mānuka, snapper and trevally. When looking at the numbers of high-quality SNPs (PHR and NMH), this array had conversion rates of 58%, 56%, 61% and 58% for each genus, respectively (S8 Table). These proportions are similar or higher than those of other plant Axiom SNP arrays designed [15,47–50], and similar or slightly lower than for other aquaculture species [16,23]. The new SNP array developed in this study is a necessary genotyping tool for these increasingly important species, for which so far researchers have had to rely on limited genetic resources.

### *Rubus* spp

High-density genotyping in raspberry and blackberry has been carried out mainly via GBS so far [27–29,31,51,52], and the development of genetic maps and QTL identification studies have been hampered by the lack of appropriate genetic resources in these species [32]. A more reliable target capture approach was used for a phylogenetic study in *Rubus* [26]; however, this method is not adapted for high-throughput genotyping and only 94 accessions were screened. The SNP array developed here, with its ∼7,000 robust SNPs validated in a diploid dataset, provides a useful tool for the routine genotyping of a large number of samples, necessary for the characterization of germplasm collections and parentage analysis in breeding programmes. Additionally, the 6,885 robust SNPs are sufficient for the construction of genetic maps and for QTL mapping analysis, and probably represent an improvement from the high-error and high-missing rate of GBS datasets. This SNP array was also tested on tetraploid *Rubus* spp. samples, and dosage genotypes could successfully be called for 4,388 out of the 12,723 markers. The structure identified in the PCA for the tetraploid samples was well explained by the collection of origin (Fig 2), suggesting that the genotype calling in this subset of SNPs was correct.

### Mānuka

Koot *et al*. [34] used pooled genome re-sequencing to study the genetic structure of mānuka collected across New Zealand. The SNP array data produced here for a subset of the populations described in Koot *et al*. [34] agrees with the expected clustering whereby NNI, ECNI and CSNI group separately. However, the two South Island provenances SWSI and NESI identified by Koot *et al*. [34] clustered together in this study (SI), and this is likely to be a result of the bias within the SNP filtering parameters, with more SNPs identifying samples from the South Island than from their individual provenances. Where comparable, the *F*_*ST*_ values obtained by both approaches were similar. These findings indicate that the SNP array can be used for reproducible genotyping across mānuka provenances. As for *Rubus* spp., medium-throughput genotyping of mānuka was previously achieved using GBS [40], which required extensive data curation [4], while the data generated with the SNP array could be directly employed in genetic diversity analyses.

### Snapper

The SNP array was able to discriminate among the different breeding populations of the snapper aquacultured strains, even though these samples were characterised by an overall high degree of relatedness, which reduced the degree of intra-specific genetic diversity. Nonetheless, the SNP data clustered the breeding populations as expected, mirroring the known relatedness among the sampled individuals (Figs 4A and B). Importantly, the SNP array performed well when used to reconstruct the pedigree of broodstock individuals, with > 90% of the offspring successfully assigned to a parental duo. The remaining offspring that could not be assigned to specific parents might derive from individuals that were not genotyped (e.g. because they died before sample collection, or because they failed at genotyping). Assignment tests of parents in the broodstock elite lines to offspring indicated skewed parental contributions to the next generation (Fig 4C), suggesting differing rates of spawning or egg generation, which is supported by previous work applying GBS data [35]. Taken together, these results support the accuracy of the SNP markers in snapper and their usefulness for genomic analysis in selective breeding programmes.

### Trevally

The SNP array data showed a clear genetic separation between the trevally sampled in Australia and those caught in New Zealand waters (Fig 5). This pattern of wide ranging panmixia is expected for many marine species with high population sizes and high dispersal [53], and has been documented for other marine teleost species in New Zealand previously (e.g. [54–56]). Interestingly, the F_ST_ values between the two clusters were unexpectedly high, denoting pronounced genetic divergence between the two clusters. This could indicate that trevally in New Zealand and Australia have been geographically isolated for prolonged periods of time and evolved rapidly during this time, possibly because of adaptive pressures to cope with different environmental gradients. Another scenario is that the Australian fishes have been misidentified and they belong to another described or even unknown carangid species in that region. As for the other species reported in this array, GBS was previously used for high-throughput genotyping in trevally [37,57], hence the ease of use of this SNP array and the accuracy of the data represent a step forward in the genomic-assisted selection and conservation for this promising aquaculture species.

### Immediate applications of the SNP array data

The SNP validation analyses carried out in this study showed the necessity of accurate genomics tools for evaluating genetic diversity and reconstructing pedigrees for applications in both breeding and conservation practices. For each of the species included in the SNP array, this study highlighted the potential of this resource in solving some of the issues encountered, as well as answering some new research questions.

#### *Rubus* spp

The *Rubus* data analysis presented some challenges linked to incorrect historical records of ploidy and taxonomy, which this new SNP array might help clarify. *Rubus* species have been reported with ploidies ranging from 2x to 12x, and the presence of aneuploids is common [58–60]. Hence, as a first step samples needed to be separated by ploidy levels to be appropriately analysed. For most of the samples evaluated here, ploidy was estimated from their assumed parentage. However, given the ease of hybridization among *Rubus* species, even between individuals with different ploidy levels, these estimates have a high error rate. Flow cytometry is often used to confirm ploidy; however, unusual cytotypes have been observed in *Rubus* at the NCGR [60], in agreement with the known occurrence of aneuploidy. Therefore, only a subset of samples from PFR and NCGR that were known to be diploid and tetraploid were analysed here. Although only robust *Rubus* SNPs were used for PCA and DAPC analyses, clustering did not match the taxonomy of the samples. This may indicate that some samples were not true-to-type, and/or that their taxonomic identification was not accurate. *Rubus* plants easily self-propagate through underground stolons, which are able to travel far from the mother plant, making their management in the orchard very difficult. Indeed, several genotypes planted next to each other in the PFR orchard proved to be identical. Furthermore, phylogenetic analysis in *Rubus* is challenging because of the common wide inter-species hybridization, apomixis, varying ploidy levels, and remarkable morphologic diversity [61]. Currently, more than 500 species are estimated to belong to this genus. Hence, many accessions at the PFR and NCGR germplasm collections have probably been misclassified taxonomically. This SNP array could be extremely helpful in revealing and/or verifying the identity of *Rubus* accessions conserved at PFR and the NCGR, as well as their degrees of relatedness, and in some cases it could even assist in their taxonomic re-classification, as it was shown in a similar study in *Pyrus* [10]. These are essential analyses for an efficient conservation programme, as well as for parental selection for cultivar improvement.

### Mānuka

This new SNP array is a fundamental resource for a more efficient management and breeding of mānuka populations. *L. scoparium* is a species of ecological, economic and cultural importance in Aotearoa-New Zealand, where it is considered taonga (treasure) by the indigenous Māori people [38]. Population genetic studies have already revealed a strong geographically dependent structure in mānuka in New Zealand [34], and further insights into its genetic diversity could enable an understanding of its adaptability to different environments, as well as assist in management practices. The possibility to perform genetic screening of mānuka populations easily and effectively will also help in the identification of loci linked to key traits for both mānuka conservation and breeding purposes, e.g. resistance to the tree-killing myrtle rust disease [62] and increased content of health-beneficial compounds in the nectar, linked to the production of high-value honey [63].

### Snapper

In snapper and many other bream species used in aquaculture (e.g. gilthead seabream and red seabream) reproduction occurs via mass spawning of selected broodstock lines, and subsequently fertilised eggs are collected and reared for on-growing. This means that parental relationships are unknown in these species, hence genetic tools are necessary for offspring parental assignment. Additionally, high-throughput genotyping with this SNP array will enable quantitative genetic studies, QTL mapping, as well as the estimate of breeding values and inbreeding rates. Routine genetic screening of new wild broodstock lines introduced in captivity, as well as the offspring generated, would also help maintain high genetic diversity in the breeding programme and maximize genetic gain.

### Trevally

The SNP array data generated in this study allowed the discrimination of two different wild trevally populations from Australia and New Zealand. Even though further work (e.g. mtDNA sequencing) is necessary to confidently resolve whether these represent two distantly related trevally populations or that a different species was instead caught in Australia, this finding indicates that the SNP array can assist in both wild population management and fisheries assessment. Furthermore, application of the SNP array to inform the selective breeding of trevally [37,57], as demonstrated for snapper, holds immense future potential, e.g. to infer relatedness of broodstock lines and to inform the selection of suitable outbred parents for new lines.

### The importance of sample quality for successful SNP array genotyping

The Axiom Analysis Suite software provides two parameters to assess sample quality: the DQC, which is based on intensities of non-polymorphic probes (i.e., that do not vary in sequence from one individual to the next) and is expected to be close to 1 for high-quality samples; and the QC call rate, which is the genotyping call rate across a subset of arrayed SNPs selected by Thermo Fisher Scientific. With no previous indication about the quality of the arrayed SNPs, as it is normal in new designs like in this study, the QC call rate was not very reliable in determining sample quality. When looking at the DQC, differences in sample quality were observed between plant and fish samples. A greater failure rate was reported in both *Rubus* and mānuka than in snapper and trevally, which had higher and more uniform DQC values (Fig 6A). There were no evident species-specific differences for QC call rate (Fig 6B), consistently with the lower reliability of this parameter in this particular case; however, in diploid *Rubus* and mānuka a number of low quality samples were identified that fell between genotype clusters and caused errors in the genotypic calls (S2 Figure). A similar behaviour was not observed in snapper and trevally. Overall, these results suggest that the plant samples had lower quality than the fish. Since all DNA samples were quantified and normalized, it is possible that these differences were caused by DNA quality. Although the quality of the DNA samples was not evaluated, for practical reasons, it is well known that DNA extraction from perennial plants is difficult, particularly because of the presence of polysaccharides and secondary metabolites [64,65]. Often DNA extraction protocols need to be optimized for each species, and scalability to large numbers of samples is particularly difficult to implement. In *Rubus*, for example, CTAB-based extractions (e.g. the Kobayashi method [66], or the protocol reported by Porebski *et al*. [67]) are usually recommended over commercial kits; however, these are difficult to implement for high-throughput 96-well plate-based extractions. In mānuka, where samples were collected in remote locations, leaf tissues were kept at room temperature and that may have caused DNA degradation. In future studies, proper DNA quality checking will be necessary to determine if samples are suitable for SNP array genotyping, and optimization of large-scale DNA extraction protocols for recalcitrant species, such as the perennial plants *Rubus* and mānuka, might be important for a higher success rate.

### Multi-species plant-animal SNP array can be run on multiplexed DNA

To the best of our knowledge, this is the first SNP array that exploits the pooling of DNA from different species into a single reaction. Here, samples from two highly divergent orders (teleost fish and dicotyledonous plants) were combined and genotyping was successfully executed for all the species screened. A number of samples were genotyped from non-pooled reactions, allowing us to assess the effect of DNA pooling on the subsequent quality of the results. It is important to note that a much larger number of pooled samples were evaluated than non-multiplexed ones. For reasons explained above, QC call rate was not considered reliable in this study and only the DQC was used for this evaluation. Overall, samples that were not pooled had a higher success rate than those that came from a mixed reaction; however, differences were observed between fish and plant samples. In snapper and trevally the differences in DQC between pooled and non-pooled samples were significant but not as marked as in mānuka and *Rubus*, where a larger variability was also observed (Fig 7A). Interestingly there was a poor correlation between the DQC of fish and plant samples from the same reaction (Fig 7C). Together these results suggest that the low quality genotyping in some samples was more likely to be caused by species-specific factors than by the pooling itself. However, it is also possible that the pooling may have disproportionally affected plant DNA, e.g. because of lower quality compared with the fish DNA. However, pooled fish samples still had very high quality (Fig 7B), supporting the use of DNA pooling in these species to reduce genotyping costs.

### Cross-species SNP performance with Japanese red seabream and yellowtail kingfish

A small number of Japanese red seabream (n = 39) and yellowtail kingfish (n = 23) DNA samples were screened over the array. The objective was to evaluate the performance of snapper and silver trevally SNPs on two closely related species within the same family, as it has been performed successfully in other taxa [16,17,22,23]. Almost all red seabream samples were successfully genotyped with snapper SNPs (S3 Table). Evaluation of a random subset of PHR and NMH SNP cluster plots showed that they often grouped together with snapper samples (S3A and S3B Figures). Some well-clustered OTV SNPs highlighted the same pattern (S3C Figure), while others showed a clear separation between red seabream and snapper (S3D Figure), indicating that this array is useful for examining the genetic diversity between these two species. This was further confirmed by the PCA (Figs 4A and B). Additionally, while the majority of the SNPs were monomorphic within the red seabream, 3,511 high-quality polymorphic SNPs (PHR, NMH or OTV) were identified, indicating that this array could be used to also evaluate intra-species genetic diversity to a certain extent. The number of polymorphic SNPs would certainly be of great value for selective breeding programmes for red seabream [68], where parent assignments following mass spawning would be needed. Concerning kingfish, all samples had very low DQC values and could not be genotypes could not be called (Fig 6A, S3 Table). This might either be because of a large divergence between trevally and kingfish genomes or because of low DNA quality, and more samples would need to be screened to confirm this.

## Conclusions

Our study clearly demonstrates that multiplexing plant and animal SNPs on the same array and pooling DNA from two distantly related species is possible and efficient. This is a promising solution for cutting costs for high-throughput genotyping in half. The SNP array designed here had a conversion rate that ranged between 56% (mānuka) and 61% (snapper), which enabled a genotyping density suitable for application in both breeding and conservation strategies. The robust and highly polymorphic SNP markers developed here for each species might be used in the future to design higher efficiency multi-species arrays, as well as being supplemented with new markers.

## Acknowledgements

We would like to acknowledge the PFR staff who assisted with the breeding and husbandry operations for the various populations; in particular Warren Fantham, who oversees the larvae rearing of finfish, and Therese Wells, who manages the post-juvenile husbandry. We also would like to thank Toshi Foster, who helped with the *Rubus* sample collection at PFR, Chris Kirk for contributing to DNA extractions in *Rubus*, Igor Ruza, Noemie Valenza-Troubat, Tom Oosting, Christina Flammensbeck, David Ashton and Matt Wylie for help with the fish sample collection, and Charles David for providing a variant-calling dataset for snapper. Finally, we would like to thank our collaborators in the *Rubus* genetics and breeding community, who used the SNP array and provided information about the quality of genotyping: Driscoll’s (Watsonville, CA, United States), Fondazione Edmund Mach (San Michele all’Adige, Italy), the Institute of Agrifood Research and Technology (Barcelona, Spain), the James Hutton Institute (Dundee, Scotland), and the National Institute of Agricultural Botany (East Malling, United Kingdom).

## Supporting Information

**S1 Appendix. Methods for sequencing, variant calling and SNP filtering, and for SNP validation in *Rubus* spp**.

**S2 Appendix. Methods for sequencing, variant calling and SNP filtering, and for SNP validation in mānuka**.

**S3 Appendix. Methods for sequencing, variant calling and SNP filtering, and for SNP validation in snapper**.

**S3 Appendix. Methods for sequencing, variant calling and SNP filtering, and for SNP validation in trevally**.

**S1 Fig. Variant calling and single nucleotide polymorphism (SNP) filtering steps for raspberry (*Rubus* subgenus *Idaeobatus*)**.

**S2 Fig. Cluster plot evaluation in *Rubus***. Cluster plots for two *Rubus* single nucleotide polymorphism (SNP) markers classified as *PolyHighResolution*. Highlighted in pink are samples with an allele_deviation_mean > 0.85.

**S3 Fig. Cluster plot evaluation for Japanese red seabream samples**. Cluster plots for snapper single nucleotide polymorphism (SNP) markers classified as *PolyHighResolution* (A), *NoMinorHom* (B) and *OffTargetVariant* (C and D). Highlighted in pink are seabream samples, while the others are snapper samples.

**S1 Table. Raspberry (*Rubus* subgenus Idaeobatus) F**_**1**_ **populations sequenced via genotyping-by-sequencing and used for variant calling**.

**S2 Table. Blackberry (*Rubus* subgenus *Rubus*) accessions re-sequenced for variant calling**. For each genotype, the repository of origin, the ploidy and the total paired-end reads are reported.

**S3 Table. Samples screened with the multi-species plant animal single nucleotide polymorphism (SNP) array**. For each sample, the genotyping batch, the plant and animal species pooled and the relative DQC and QC call rate values are reported.

**S4 Table. Mānuka samples used for single nucleotide polymorphism (SNP) validation**. For each sample, the region of origin is reported. A table exhibiting the number of samples per provenance is also reported.

**S5 Table. Trevally samples used for single nucleotide polymorphism (SNP) validation**. For each sample, the country of harvesting is reported. A table exhibiting the number of samples per country of origin is also reported.

**S6 Table. Classes of mānuka single nucleotide polymorphisms (SNPs) identified based on their minor allele frequency (MAF) values in each or a combination of gene pools**.

**S7 Table. Details of the single nucleotide polymorphism (SNP) markers included in the multi-species 60K SNP array**. For each SNP the position on the reference genome and the flanking sequences are reported.

**S8 Table. Number of single nucleotide polymorphism (SNP) markers by species and Axiom classification**.

