## Supplementary material for "A multiplexed plant-animal SNP array for selective breeding and species conservation applications": S1 Appendix

### ***Rubus* spp.**

#### Sequencing, variant calling and SNP filtering

**Raspberry.** Data from genotyping experiments produced using reduced-representation genotyping-by-sequencing (GBS) were available for seven F_1_ *Rubus* subgenus *Idaeobatus* populations, including sequences from the offspring and, where available, the parents and grandparents (S1 Table). One family was developed at North Carolina State University from a cross between accession NC493 (*R. parvifolius* × *R. idaeus* ‘Cherokee’) and ‘Chilliwack’ (CW) (*R. idaeus*). GBS data for NC493×CW were retrieved from Jibran *et al.* [1]. The raw VCF file was used, which included 649,597 single nucleotide polymorphisms (SNPs) for all offspring and parents with no filtering applied. Six families generated at Plant & Food Research (PFR) by crossing *R. idaeus* breeding selections consisted of: X14.102 (n = 157), X16.015 (n = 94), X16.093 (n = 47), X16.095 (n = 199), X16.109 (n = 56), and X16.111 (n = 49). For these populations, GBS libraries were prepared and sequenced using the protocol of Jibran *et al.* [1]. Reads were then trimmed of TrueSeq Illumina adapters with Trim Galore v 0.4.3 (<https://github.com/FelixKrueger/TrimGalore>), de-multiplexed with *fastq_multx* in ea-utils v 1.1.2-806 (<https://github.com/ExpressionAnalysis/ea-utils>), and aligned to the *R. occidentalis* v 3.0 genome [2] with BWA-mem v 0.7.17 [3]. Variant calling was performed with SAMtools v 1.7 (*mpileup*) and bcftools v 1.10.2 (multiallelic-caller) [4]. For the X14.102, X16.015 and NC493×CW families, INDELs were filtered out, and individuals with $>$ 80% missing rate and SNPs with RMS mapping quality $<$ 20, depth $<$ 10 or $>$ 1,000 were removed. The same parameters were used for the other families, except that SNPs were filtered for RMS mapping quality $>$ 20, depth $>$ 8, and minor allele frequency (MAF) $>$ 0.05.

All the above datasets were merged with VCFtools v 0.1.14 [5] *vcf-merge*, to identify and eliminate sites with another SNP within 30 bp up- or down-stream (thinning). The list of remaining SNPs was intersected back with each of the four datasets, from which a core of “validated SNPs” was subsequently identified. “Validated SNPs” had no inconsistencies between technical replicates of the same genotype, exhibited $<$ 5% Mendelian errors, $<$ 80% missing data and were polymorphic within the families; additionally, individuals with $>$ 10% error rate were also removed. Finally, genotypic data for the families X16.093, X16.095, X16.109, X16.111 and NC493×CW were imported into JoinMap v5.0 [6] and grouped into Linkage Groups (LGs) with a LOD $\geq$ 10; SNPs that were not successfully included into one of the expected seven LGs were eliminated from the “validated SNPs” datasets of those families. As a last filtering step, A/T and C/G SNPs were discarded and a unique list of SNPs from all datasets was obtained. Finally, a total of 859 SNPs, designed on candidate genes controlling sugar content and validated with a KASP assay [7], were added to the merged dataset.

A visual representation of the variant calling and filtering performed in *Rubus* is reported in S1 Fig.

**Blackberry.** Whole-genome sequencing (WGS) data for 27 blackberry cultivars and advanced selections from the University of Arkansas System Division of Agriculture (UArk) and Agricultural Research Service of United States Department of Agriculture (USDA-ARS) breeding programmes (S2 Table) were used for SNP selection. Samples were sequenced on Illumina HiSeq 2500 to generate from 84.6 to 123.7 million 2×150 bp paired-end reads per sample. Raw Illumina reads were processed to remove contaminating sequencing adapters and low-quality reads using the CLC Genomics Workbench (Qiagen, Hilden, Germany). Adapter-trimmed, high-quality reads were then mapped to a contig-scale genome assembly of the diploid blackberry ‘Hillquist’ (*R. argutus*) [8]. Mapping was performed with 90% identity and 90% read coverage parameters using CLC Genomics Workbench V20.0. The mapped bam file was sorted using Samtools and the duplicate reads were marked using sambamba [9]. Variant calling was performed on the de-duped bam file using Freebayes version 1.3.2 [10]. The Freebayes output was further filtered using bcftools and custom scripts to obtain high-quality and biologically-relevant SNP markers. Only biallelic SNPs (no A/T and C/G) with at least one homozygous individual in the panel, less than 33% missing data, and no other flanking variants in a 30 bp up- or down-stream window were selected. Finally, these SNPs were re-aligned to a new chromosome-length assembly of *R. argutus* [11] and only those that mapped to a unique position were selected.

The flanking sequences of the blackberry SNP were BLAST searched against the *R. occidentalis* genome and those of the raspberry SNPs were BLAST searched against the *R. argutus* genome [11], with the objective of identifying overlapping SNPs between the two datasets and retaining one. In addition, blackberry SNPs with multiple hits on the raspberry reference genome were discarded, as they could result in erroneous genotypic calls in raspberry individuals.

#### SNP validation

Diploid and tetraploid *Rubus* samples from PFR, (USDA-ARS) - National Clonal Germplasm Repository (NCGR), and UArk were analysed.

**Diploids.** A total of 477 samples were estimated to be diploid, including 332 from PFR, 143 from NCGR, and two from UArk. These samples were analysed together in the Axiom Analysis Suite software. Cluster plots of a random subset of *PolyHighResolution* SNPs (PHR) were observed to verify the quality of the call and make necessary adjustments. Samples with an “allele_deviation_mean” value (i.e. the average of the absolute difference between the log_2_ allele signal estimate and its median across all SNPs) higher than 0.85 were then removed and the remaining samples were re-analysed. SNPs categorized as PHR and *NoMinorHom* (NMH) were then verified for consistency in biologically replicated samples (i.e. samples collected twice or more from the same plant). The similarity of duplicated samples was checked by calculating pairwise identity-by-state (IBS) values in the R package SNPRelate v1.18.1 [12]. All replicates with an IBS $>$ 0.97 were considered truly identical and used to remove markers with inconsistent genotypic calls and identify a subset of robust SNPs to use for follow-up analysis. A Principal Component Analysis (PCA) was run on the diploid good-quality samples using the robust SNPs filtered for MAF $>$ 0.05 and LD-pruned with a threshold of 0.2. Additionally, a Discriminant Analysis of Principal Components (DAPC) was run in the R package adegenet v2.1.2 [13–15].

**Tetraploids.** The overall number of tetraploid samples analysed was 739, including 262 from PFR, 66 from NCGR and 411 from UArk. Summarized signal intensities for all 12,723 *Rubus* SNPs for these samples were obtained in the Axiom Analysis Suite software and then imported into R for dosage calling with the package fitPoly v3.0.0 [16,17]. The command *saveMarkerModels* was used with a p-value threshold of 0.9. The dataset was subsequently filtered for missing rate, applying a 20% cut off for both samples and SNPs. Finally, a PCA was run using the R package polymapR v1.1.2 [18].
