## Supplementary material for "A multiplexed plant-animal SNP array for selective breeding and species conservation applications": S2 Appendix

### **Mānuka**

#### Sequencing, variant calling and SNP filtering

The pooled sequencing data from Koot *et al.* [1] were used for SNP selection. This dataset consisted of 68 pooled populations of mānuka sampled across New Zealand. Five gene pools were identified amongst these populations by Koot *et al*. [1], namely in the Northern North Island (NNI), the Central and Southern North Island (CNI), the East Coast North Island (ECNI), and two gene pools in the South Island representing the North-East (NESI) and South-West of the South Island (SWSI), respectively. The variants called in the 68 pooled populations were filtered using VCFtools, by keeping SNPs with MAF $>$ 0.05, a mean DP of 100, and no missing data and discarding A/T and C/G SNPs. Additionally, MAFs were calculated and averaged across populations within each gene pool and used to identify SNPs specific to each of the gene pools, as well as to the North and South Island respectively. The SNP locations were based on the reference genome of *Leptospermum scoparium* ‘Crimson Glory’ [2].

#### SNP validation

A subset of 264 samples used for pool sequencing and variant detection by Koot *et al.* [1] was employed for SNP validation. The samples were chosen as representatives of five gene pools: NNI, ECNI, CSNI, NESI and SWSI (S4 Table). Population structure was investigated using K-means clustering and DAPC analyses in the R package adegenet. Weir and Cockerham's pairwise *F_ST_* distances [3] and accompanying p‐values were estimated among gene pools using the R package StAMPP v1.6.3 [4] and applying nboots = 1000, percentage = 95.
