## Supplementary material for "A multiplexed plant-animal SNP array for selective breeding and species conservation applications": S3 Appendix

### **Snapper**

#### Sequencing, variant calling and SNP filtering

Both parental individuals and six offspring of ten F_1_ families from the PFR snapper breeding programme, accounting for a total of 80 samples, were sequenced using the Illumina Novaseq technology. Reads were trimmed using Trimmomatic v0.36 [1], then aligned to the *Chrysophrys auratus* v 1.0 male reference genome [2] with BWA-mem. Variants were called with three different software (SAMtools mpileup and bcftools multiallelic-caller; GATK v4.0.3.0 HaplotypeCaller [3]; and FreeBayes v1.3.1) and combined into a final consensus set. SNPs with another polymorphism within 30 bp up- or down-stream were removed, as well as those with $>$ 20% missing data, DP $>$ 6,493 ($=$ average DP $+$ 3 standard deviations), and MAF $<$ 0.05. Multi-allelic and A/T and C/G SNPs were also discarded. Finally, the remaining SNPs were pruned for linkage disequilibrium (LD) using bcftools +prune, retaining only four SNPs with r^2^ $>$ 0.85 per 100 kb window.

#### SNP validation

Genotypes for snapper were called keeping the two batches separated, as a large number of samples were screened (n $=$ 2,525 and n $=$ 1,719, respectively for first and second batch). Seabream samples (n $=$ 39) were included in the first batch. The two datasets were then merged and only SNPs classified as PHR in both datasets were kept for subsequent analysis. A PCA was run using the R package SNPRelate to examine the genetic diversity between snapper and seabream, as well as the structure of the snapper samples. Additionally, pedigree reconstruction was carried out for a subset of snapper samples corresponding to three separate broodstock lines generated at PFR (Broodstock 1: 29 parents, 1,114 offspring; Broodstock 2: 35 parents, 965 offspring; Broodstock 3: 54 parents, 1,153 offspring). The KING-robust algorithm [4] in SNPRelate was used to calculate pairwise kinship coefficients (*k*) and IBD0 values between samples. Putative trios were identified as those with the highest ratio of *k*:IBD0 between parent and offspring samples. True trios were then confirmed if they had less than 2% Mendel errors as calculated using the python package scikit-allel v1.3.3 (https://github.com/cggh/scikit-allel).
