## Supplementary material for "A multiplexed plant-animal SNP array for selective breeding and species conservation applications": S4 Appendix

### **Trevally**

#### Sequencing, variant calling and SNP filtering

The dataset employed here included 17,795,808 SNPs that resulted from the calling and quality-filtering performed by Valenza-Troubat *et al.* [1] on WGS reads of 13 trevally samples, representing parental individuals of the PFR breeding programme, and using the male reference genome for trevally [2]. As for snapper, this dataset was filtered by removing SNPs that had another polymorphism within 30 bp up- or down-stream, $>$ 20% missing data, DP $>$ 445 ($=$ average DP $+$ 3 standard deviations), MAF $<$ 0.05, and were multi-allelic or A/T or C/G. The remaining SNPs were then LD-pruned, keeping only five SNPs with r^2^ $>$ 0.85 per 100 kb window.

#### SNP validation

A subset of 978 trevally samples, representing samples from fish caught in the wild in New Zealand and Australia, was analysed for SNP validation (S5 Table). SNPs that were classified as PHR and NMH were filtered for MAF $>$ 0.05 and LD-pruned with a threshold of 0.2. A PCA was run using the *prcomp* function in R (setting *center = TRUE*, *scale. = TRUE*) and results were plotted with the factoextra package v1.0.7 [3]. Weir and Cockerham's weighted *F_ST_* was estimated for country of origin in SNPRelate.
