## Supplementary figures and images for "A multiplexed plant-animal SNP array for selective breeding and species conservation applications"

### S1 Figure

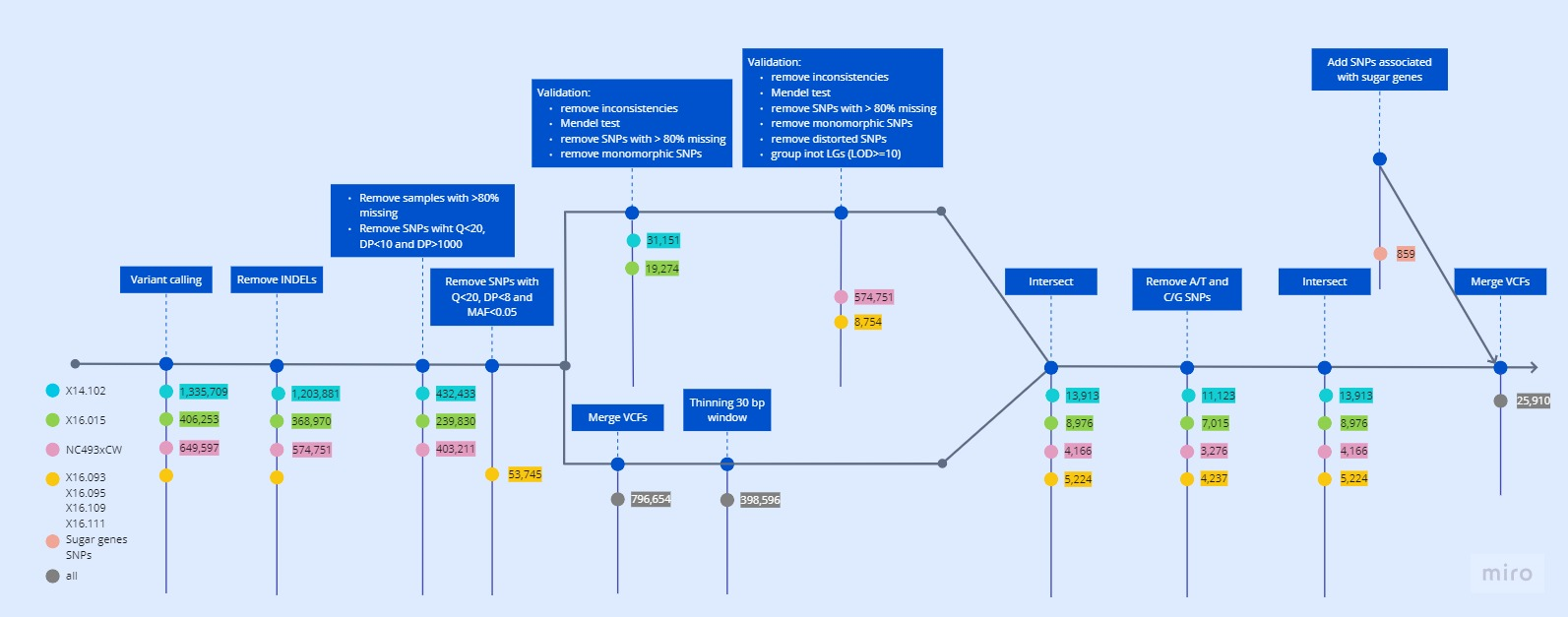

### S2 Figure

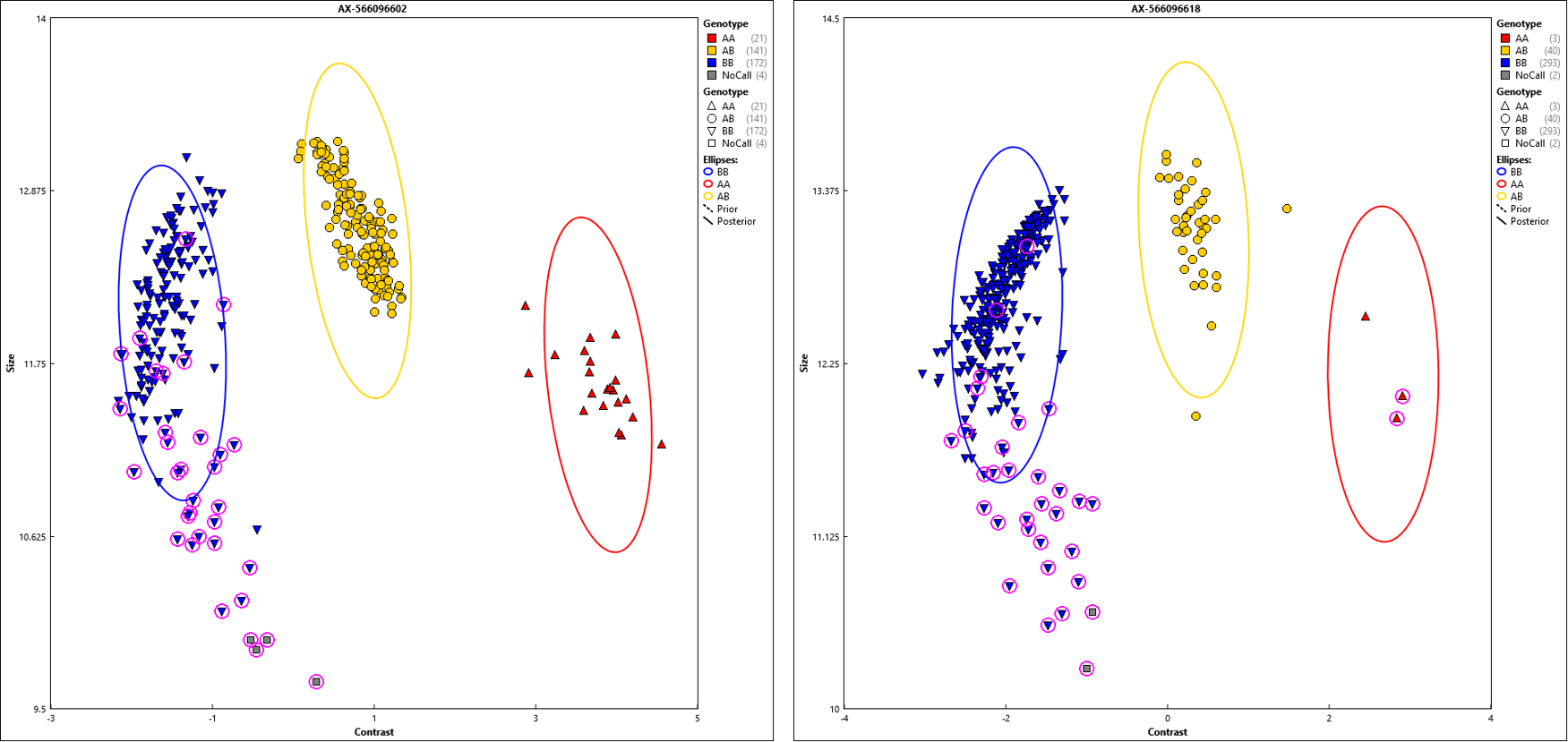

### S3 Figure

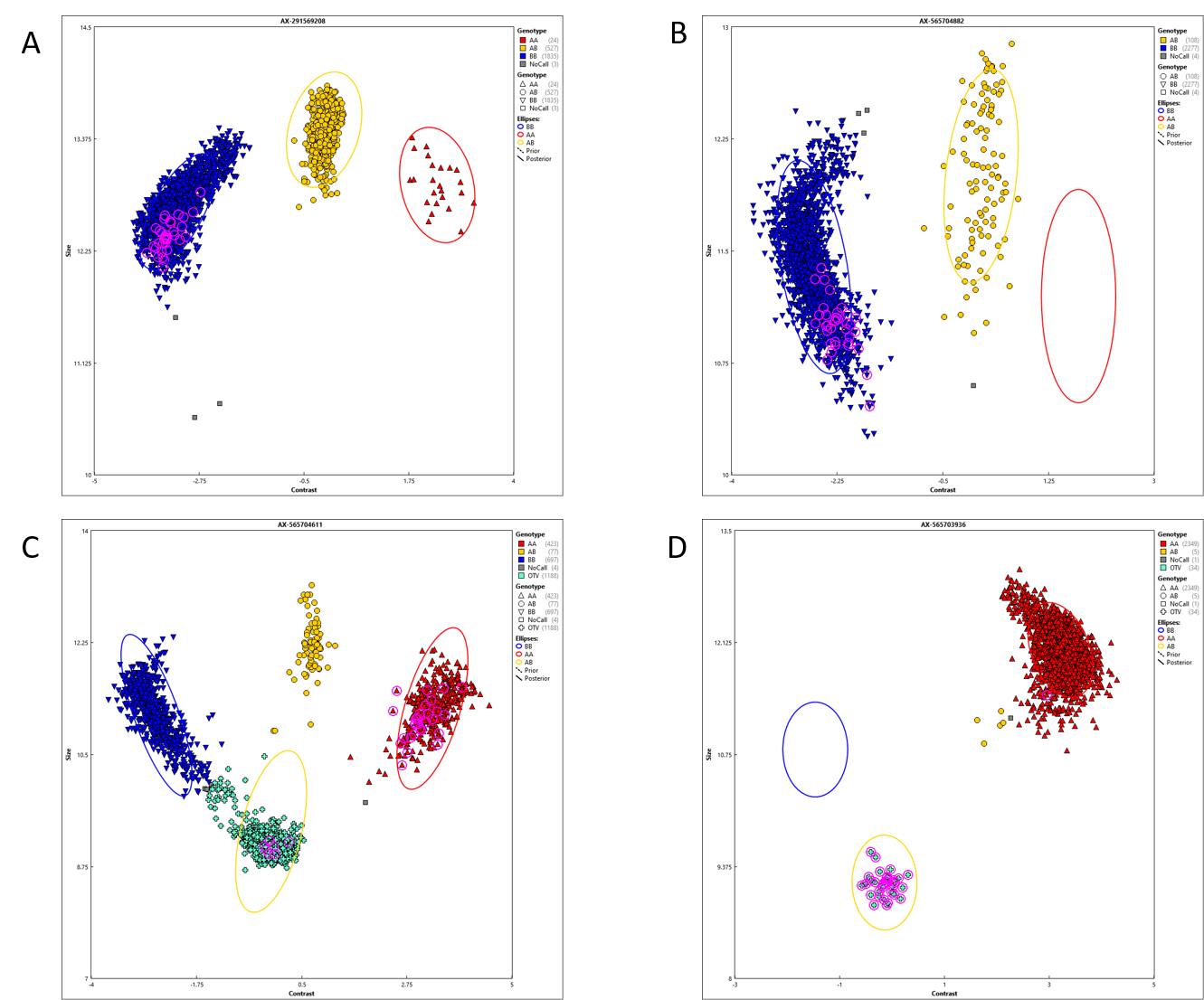
